# Effect of Phosphorylation on the Collision Cross Sections of Peptide Ions in Ion Mobility Spectrometry

**DOI:** 10.1101/2020.06.15.151639

**Authors:** Kosuke Ogata, Chih-Hsiang Chang, Yasushi Ishihama

## Abstract

The insertion of ion mobility spectrometry (IMS) between LC and MS can improve peptide identification in both proteomics and phosphoproteomics by providing structural information that is complementary to LC and MS, because IMS separates ions on the basis of differences in their shapes and charge states. However, it is necessary to know how phosphate groups affect the peptide collision cross sections (CCS) in order to accurately predict phosphopeptide CCS values and to maximize the usefulness of IMS. In this work, we systematically characterized the CCS values of 4,433 pairs of mono-phosphopeptide and corresponding unphosphorylated peptide ions using trapped ion mobility spectrometry (TIMS). Nearly one-third of the mono-phosphopeptide ions evaluated here showed smaller CCS values than their unphosphorylated counterparts, even though phosphorylation results in a mass increase of 80 Da. Significant changes of CCS upon phosphorylation occurred mainly in structurally extended peptides with large numbers of basic groups, possibly reflecting intramolecular interactions between phosphate and basic groups.

## Introduction

Protein phosphorylation is a reversible post-translational modification that influences protein folding, activity, protein-protein interaction and subcellular localization^1^. It is well known to play key roles in intracellular signal transduction pathways regulating numerous cell functions^2^. Various techniques have been developed to monitor altered phosphorylation events in cells, and mass spectrometry is currently one of the most powerful techniques for proteome-wide experiments^3,4^. In LC/MS-based shotgun proteomics, MS/MS spectra are mainly used to identify peptides but other information, such as LC retention time, can also help to increase the confidence of sequence assignments^5^. With recent improvements in prediction models^6–10^, peptide retention time information has become an increasingly powerful aid for peptide identification and quantitation, especially in target mode^11,12^ and data-independent acquisition mode^13^ analyses. However, the models for predicting the retention times of phosphopeptides are not yet mature, in part due to the absence of a comprehensive understanding of the retention mechanisms of the phosphopeptides. Accordingly, the retention mechanism of phosphorylated peptides has been investigated by comparing pairs of phosphorylated and unphosphorylated peptides^14–17^.

The insertion of ion mobility spectrometry (IMS) between LC and MS has recently attracted interest as a means of improving peptide identification in both proteomic and phosphoproteomic experiments^18–22^. IMS adds a dimension of separation and provides structural information that is complementary to LC and MS, because it separates ions on the basis of differences in their shapes and charge states. Again, though, it is necessary to know how phosphate groups affect the peptide collision cross sections (CCS) in order to accurately predict phosphopeptide CCS values and to maximize the usefulness of IMS. Previous IMS-MS studies^23–26^ on small numbers of singly and doubly charged phosphopeptide ions have indicated that the CCS of phosphopeptides is smaller than that of unmodified peptides of equivalent mass. Based on these findings, the model using the intrinsic size parameter (ISP)^27,28^ for the prediction of CCS of phosphopeptides has been extended ^24^; the ISP provides the average contribution of each amino acid residue or modification to the CCS of a peptide, and can predict the CCS based on the amino acid composition of the peptide. However, this model does not currently take into account sequence-specific features, such as positional information or residue combinations. Also, in this model, changes in CCS upon phosphorylation are treated as increases in CCS^24,29^, and it is not possible to explain the decrease in CCS due to phosphorylation that is seen in certain sequences^24,30^. Furthermore, the model was developed using doubly charged peptides with molecular weights ranging from 1,000 Da to 1,400 Da, and thus cannot be applied directly to larger, more highly charged ions in the 600 Da to 5,000 Da range that are commonly found in proteomics experiments. Therefore, further data collection is needed to establish an improved model for CCS prediction of phosphopeptides.

Here, we systematically characterized the CCS values of phosphopeptides and their corresponding unmodified counterparts using trapped ion mobility spectrometry (TIMS). The CCS values of 6,544 phosphopeptide ions from HeLa tryptic digests were examined, yielding 4,433 CCS pairs of phosphopeptide and unphosphorylated peptide ions with charge states ranging from 2+ to 4+ in the mass range from 800 Da to 4500 Da. Our results show that large changes in CCS occur mainly in structurally extended peptides with multiple basic groups.

## Materials and Methods

### Materials

UltraPure™ Tris Buffer was purchased from Thermo Fisher Scientific (Waltham, MA, USA). Sequencing grade modified trypsin was purchased from Promega (Madison, WI, USA). Water was purified by a Millipore Milli-Q system (Bedford, MA, USA). Porous titanium dioxide beads (TitansphereTiO, 10 μm) and Empore disks (SDB-XC and C8) were obtained from GL Sciences (Tokyo, Japan). All other chemicals and reagents were purchased from Fujifilm Wako Pure Chemical (Osaka, Japan), unless otherwise specified.

### Cell culture and protein digestion

HeLa cells were cultured to 80% confluence in DMEM containing 10% FBS in 10 cm diameter dishes. Cells were washed twice with ice-cold PBS, collected using a cell scraper, and pelleted by centrifugation. Hela cell lysates were digested by means of the phase-transfer surfactant (PTS)-aided trypsin digestion protocol, as described previously^31^. After digestion, the sample was desalted using SDB-XC StageTips^32^.

### Phosphopeptide enrichment

Metal oxide chromatographic (MOC) tips were prepared as described previously^33^. Briefly, C8 StageTips packed with TiO2 beads (0.5 mg/tip) were equilibrated with 80% ACN with 0.1% trifluoroacetic acid (TFA) and 300 mg/mL lactic acid as a selectivity enhancer (solution A). The samples were diluted with an equal amount of solution A and loaded onto the MOC tips. After washes with solution A and 80% ACN with 0.1% TFA, phosphopeptides were eluted with 0.5% piperidine. The eluate was acidified with 10% TFA and desalted using StageTips.

### Dephosphorylation of phosphopeptides

One-third of the phosphopeptide sample was dried and dissolved in 25 μL of 100 mM Tris-HCl buffer (pH 9.0). Alkaline phosphatase (from calf intestine; 5 units) was added, and the solution was incubated for 3 hours at 37°C. After the reaction, the buffer was acidified by adding 10% TFA 10 μL. The samples were desalted using StageTips.

### HPLC fractionation

The phosphopeptides and their dephosphorylated counterparts were mixed, dissolved in 4% ACN with 0.5% TFA and injected onto a Protein-RP column (2.0 mm i.d., 150 mm length, 5 μm C4-silica (USP L26), 20 nm pore) (YMC, Kyoto, Japan) using an ACQUITY UPLC H-Class Bio system (Waters, Milford, MA, USA) with a UV detector (280 nm). The injection volume was 10 µL and the flow rate was 200 µL/min. Peptides were separated with a two-step linear gradient of 4-40% ACN in 30 min, 40-80% ACN in 1 min and 80% ACN for 4 min with acetic acid as an ion-pair reagent.

### LC/TIMS/Q/TOF analysis

NanoLC/TIMS/Q/TOF analyses were performed on a timsTOF Pro (Bruker, Bremen, Germany) connected to an Ultimate 3000 pump (Thermo Fisher Scientific) and an HTC-PAL autosampler (CTC Analytics, Zwingen, Switzerland). Peptides were separated on self-pulled needle columns (150 mm length, 100 μm ID, 6 μm needle opening) packed with Reprosil-Pur 120 C18-AQ 3 μm reversed-phase material (Dr. Maisch, Ammerbuch, Germany). The injection volume was 5 μL, and the flow rate was 500 nL/min. Separation was achieved by applying a linear gradient of 4−32% ACN in 50 min with 0.5% acetic acid as an ion-pair reagent. The TIMS section was operated with a 200 ms ramp time and a scan range of 0.6-1.5 Vs cm^-2^. One cycle was composed of 1 MS scan followed by 10 PASEF MS/MS scans. MS and MS/MS spectra were recorded from *m/z* 100 to 1,700. A polygon filter was applied to avoid selection of singly charged ions. The quadrupole isolation width was set to 2 Da. The TIMS elution voltage was calibrated linearly to obtain reduced ion mobility coefficients (1/K_0_) using three selected ions (*m/z* 622, 922, and 1222) of the ESI-L Tuning Mix (Agilent, Santa Clara, CA, USA).

### Database searching and data processing

Peptides and proteins were identified through automated database searching using MaxQuant^34,35^ (version 1.6.14.0) in the TIMS-DDA mode against the human database from UniprotKB/Swiss-Prot release 2017/04 with a strict Trypsin/P specificity allowing for up to 2 missed cleavages. Carbamidomethyl (C) was set as a fixed modification. Oxidation (M), Acetyl (Protein N-term) and Phospho (STY) were allowed as variable modifications. The resulting evidence.txt file was used for the analysis.Collision cross sections were calculated from ion mobility coefficients (1/K_0_) using MaxQuant.

### Data availability

The MS raw data and analysis files have been deposited with the ProteomeXchange Consortium (http://proteomecentral.proteomexchange.org) via the jPOST partner repository^36,37^ (https://jpostdb.org) with the data set identifier PXD019746.

## Results & Discussion

First, a sample containing thousands of pairs of phosphorylated and unphosphorylated peptides was prepared by applying the workflow used to profile the differences in retention times between phosphorylated and unphosphorylated peptides^17^. After fractionation, the sample was analyzed by LC/TIMS/MS/MS to obtain paired CCS values. To simplify the analysis, peptides with multiple CCS values were aggregated to the CCS with the highest peak intensity, and peptides containing methionine oxidation and N-terminal acetylation were removed. Finally, CCS values of 6,544 phosphopeptide ions and 22,666 unphosphorylated peptide ions, including more than 4000 paired CCS values, were obtained from the HeLa tryptic digest (Table 1).

**Table 1.** Summary of identification results

|  | Phosphopeptides | Unphosphorylated peptides |
| --- | --- | --- |
| Unique peptide ions<br>(Sequence + Modification + Charge)<sup>a</sup> | 6,544 | 22,666 |
| Unique peptides<br>(Sequence + Modification)<sup>a</sup> | 5,965 | 20,298 |
| Unique peptide sequences<br>(Sequence)<sup>a</sup> | 4,942 | 20,298 |
| Unique peptide ion pairs (P / uP)<sup>b</sup><br>(Sequence + Modification + Charge)<sup>a</sup> | 4,433 |  |
| Unique peptide pairs (P / uP)<sup>b</sup><br>(Sequence + Modification)<sup>a</sup> | 4,080 |  |

a. unique key, b. P: phosphopeptides uP: unphosphorylated peptides

Figure 1A shows the CCS versus *m/z* plots obtained for unphosphorylated peptide ions. In this plot, the doubly, triply and quadruply charged populations are clearly separated. There is a strong correlation between mass and CCS within each charge state, and the triply charged species are clearly split into two subpopulations: extended and compact forms. These observations agree with previous IMS-MS findings^18,38–40^. Figure 1B shows CCS versus *m/z* plots of phosphorylated peptide ions. The trend for phosphorylated peptide ions was similar to that for unphosphorylated peptide ions. However, in the case of the triply charged species, the relative density of the smaller CCS subpopulation appeared to be higher for the phosphorylated peptide ions than for the unphosphorylated peptide ions. For further investigation, phosphorylated and unphosphorylated peptide ions were classified according to their charge state and overlaid on the CCS versus mass plot (Figure 2A-C). The difference in the average CCS values of phosphorylated and unphosphorylated peptides in each molecular weight bin, plotted in Figure 2D, shows that phosphorylated peptides are generally smaller than unmodified peptides of the same mass, which is consistent with previous IMS-MS studies^23–25^. While previous studies have focused primarily on singly and doubly charged peptide ions, we have confirmed the same trend for triply and quadruply charged peptides.

**Figure 1.**
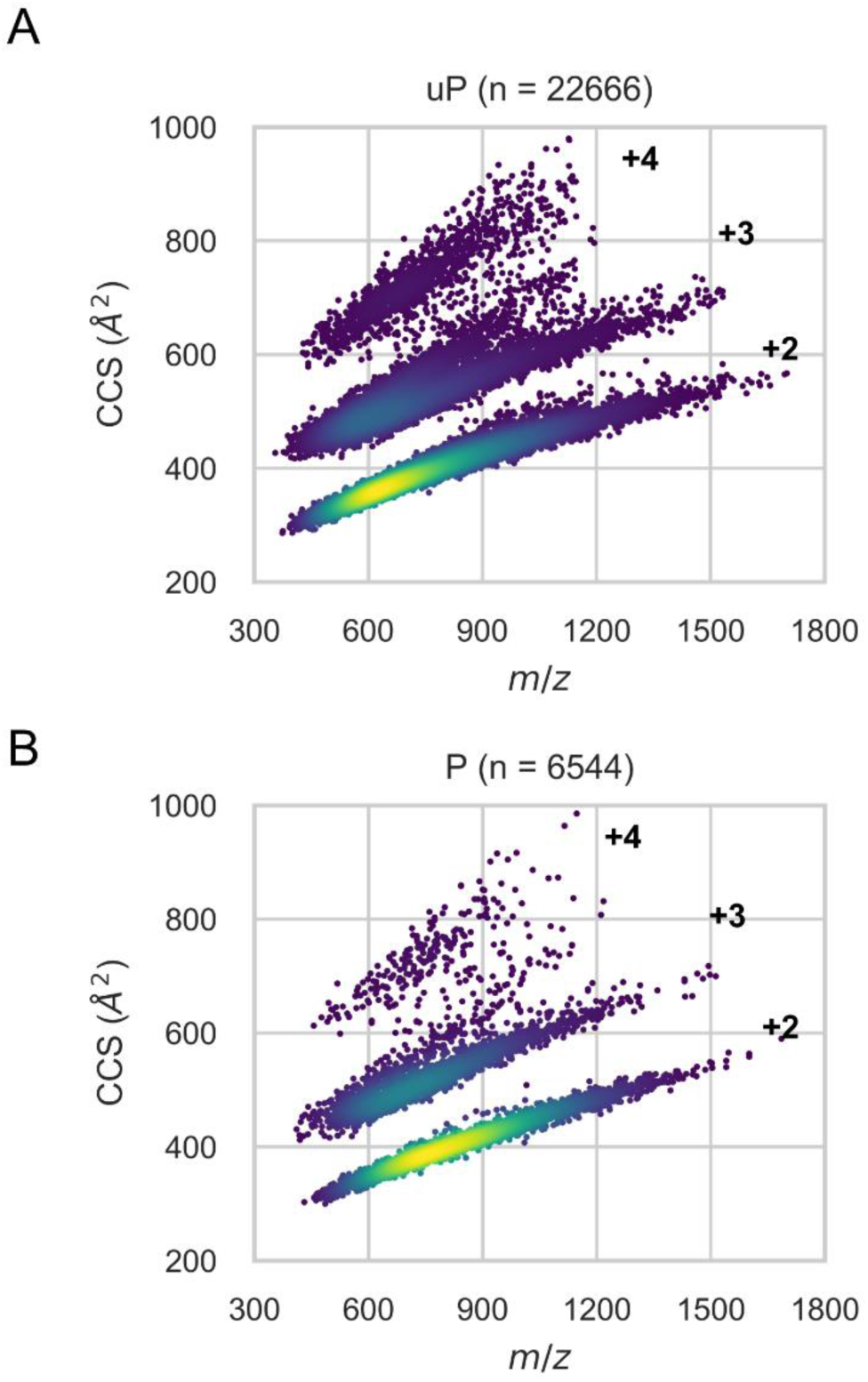
CCS for (A) unphosphorylated and (B) phosphorylated peptide ions as a function of *m/z.* The lower, middle, and upper populations consist of doubly, triply, and quadruply charged ions, respectively. The triply charged ions are further divided into two subgroups: extended ions on the upper side and compact ions on the lower side. P: phosphopeptides, uP: unphosphorylated peptides.

**Figure 2.**
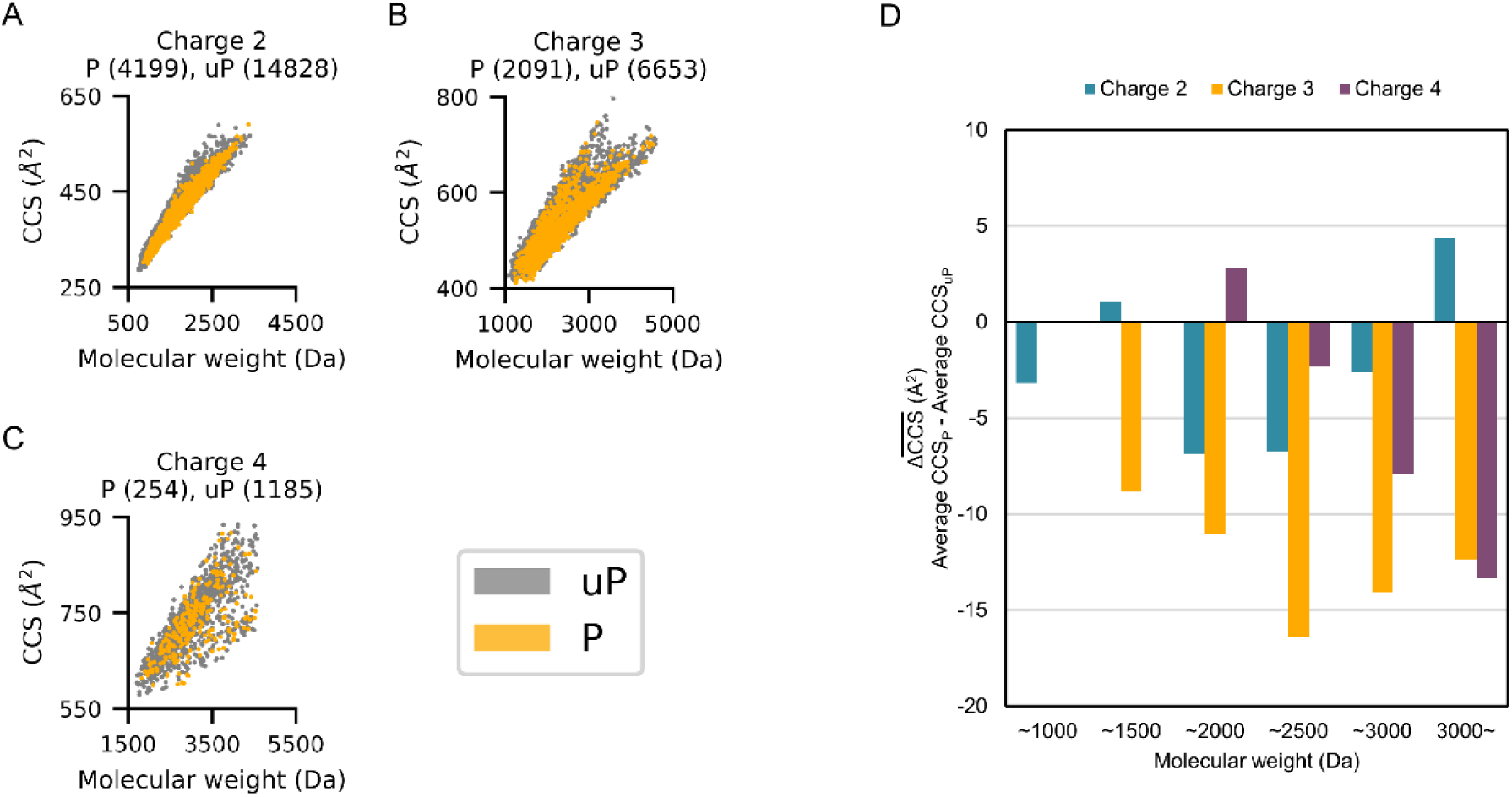
Mass versus CCS plot showing the difference between phosphorylated and unphosphorylated peptide distributions. (A,B,C) CCSs for unphosphorylated and phosphorylated peptide ions as a function of peptide mass. (D) Differences between averaged CCS values for phosphopeptides and unphosphorylated peptides in various molecular weight bins. CCS of phosphopeptides were generally smaller than those of unmodified peptides of equivalent mass. P: phosphopeptides, uP: unphosphorylated peptides.

To further evaluate the effect of phosphorylation of peptides on CCS, pairs of mono-phosphorylated and unphosphorylated peptides with identical sequences were compared. Considering the correlation between CCS and mass, the mono-phosphorylated peptide should have a larger CCS than the unmodified sequence, since phosphorylation results in an 80 Da mass increase. In fact, the median CCS of the mono-phosphopeptides was 4.0 Å^2^ larger than that of the unphosphorylated peptides (Figure 3A). However, 32.0% (1419/4433) of the mono-phosphorylated peptide ions had lower CCS values than the corresponding unphosphorylated form, suggesting that a significant number of peptides underwent compaction of their conformation upon phosphorylation (Figure 3A). The content of the CCS-compressed phosphopeptides (ΔCCS < 0) was much higher than that reported for 66 doubly charged phosphopeptides, which was 19.7% (13/66)^24^. Indeed, for doubly charged mono-phosphopeptides in our data, the content was 23.4% (642/2743), as shown in Figure 3B. In the case of triply and quadruply charged species, the contents of the phosphopeptides with ΔCCS < 0 were 46.2% (688/1488) and 44.1% (89/202), respectively (Figure 3C, D). These results can be partly explained by the molecular weight distribution of the peptides in each charge state, since the more highly charged peptides have higher molecular weights and the ΔCCS values were weakly negatively correlated with the molecular weight of the peptides (Figure 4). For larger peptides, the mass shift of 80 Da was small compared to the total mass of the peptide, and thus the effect of the mass increase on CCS was expected to be relatively small. However, for some of the larger peptides, a significant change in CCS was observed, which could be attributed to the rearrangement of peptide conformation upon phosphorylation. On the other hand, relatively small doubly charged peptides, such as those examined in the previous study, did not show such a substantial change, and mainly showed a positive shift in CCS due to the positive mass shift.

**Figure 3.**
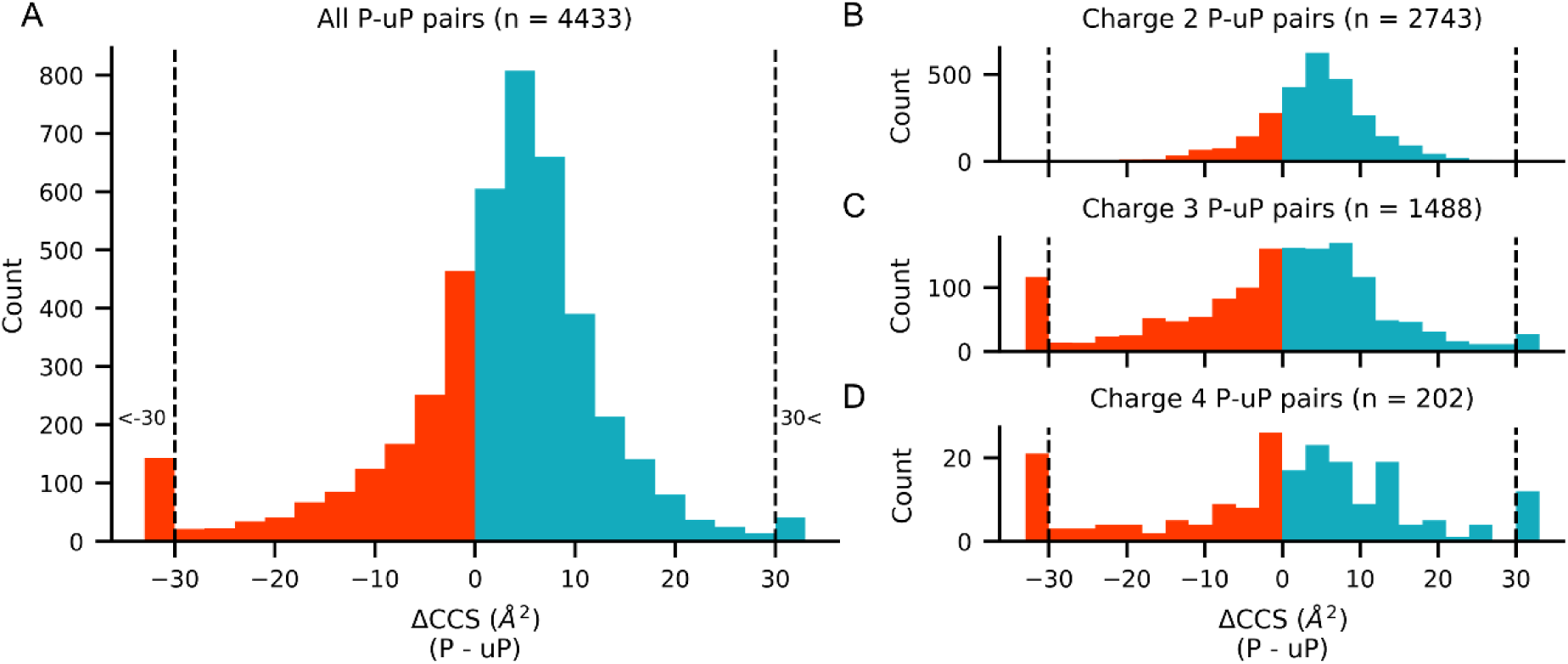
Distribution of CCS differences between phosphopeptide and unphosphorylated peptide pairs. ΔCCS was calculated by subtracting the unphosphorylated peptide CCS from the phosphopeptide CCS. ΔCCS values greater than 30 or less than -30 were aggregated. (A) Histogram of ΔCCS of all identified peptide pairs. (B-D) Histogram of ΔCCS for each charge state. P: phosphopeptides, uP: unphosphorylated peptides.

**Figure 4.**
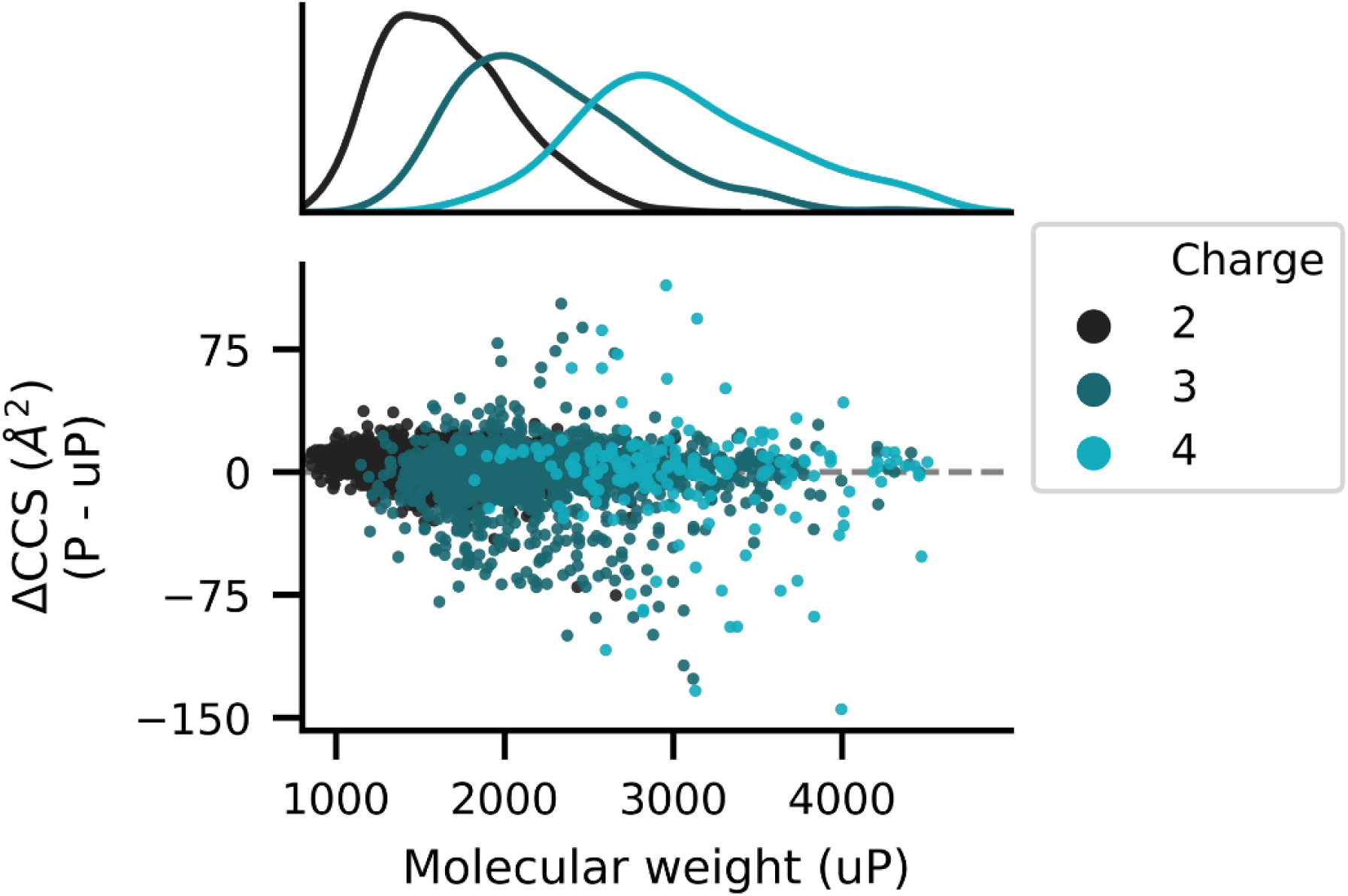
ΔCCS values for peptide ions with each charge state as a function of peptide molecular mass. The dashed line represents the ΔCCS value of zero. P: phosphopeptides, uP: unphosphorylated peptides.

However, even within the specific MW range of 1,500 Da to 2,000 Da, 26.8% (306/1140) and 48.0% (231/481) of doubly and triply charged mono-phosphorylated peptide ions, respectively, showed CCS compaction upon phosphorylation. These results suggest that ions with higher charge states show more pronounced conformational compaction upon phosphorylation. This may be attributed to the wider distribution of CCS for the triply charged ions than for the doubly charged ions, as has been indicated in previous reports^38,41^. The structural diversity of the triply charged ions may allow for a more dynamic change of structure upon phosphorylation, but further studies are needed to test this idea.

To assess what kinds of peptide features are associated with the CCS reduction upon phosphorylation, we examined the relationship between *m/z*, CCS and ΔCCS values (Figure 5). The colors of the markers indicate the ΔCCS values; the most marked change in CCS appears to be associated with the transition between extended and compact subpopulations of triply charged ions. Although less pronounced, similar trends were observed for doubly and quadruply charged ions.

**Figure 5.**
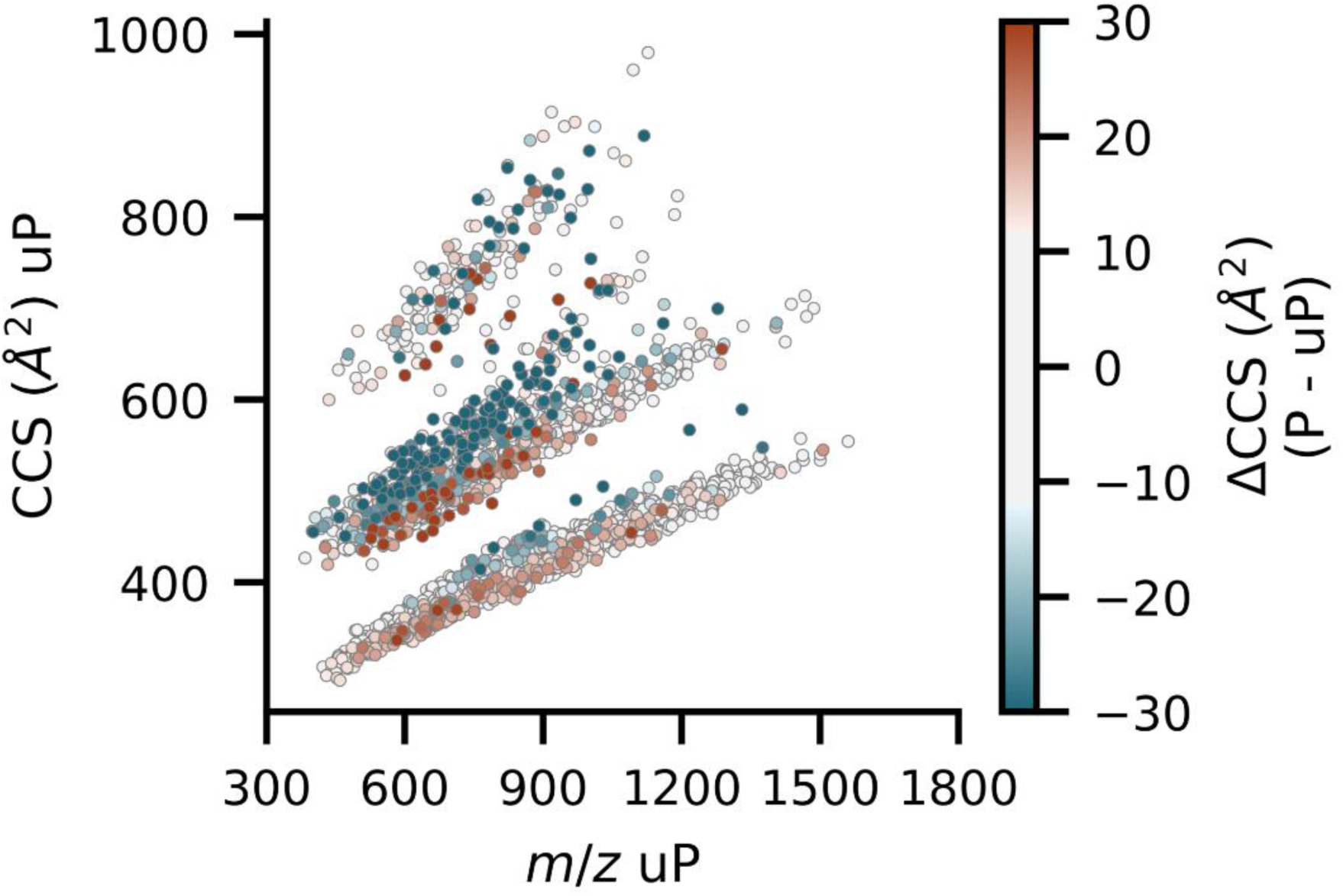
CCS for unphosphorylated peptide ions as a function of *m/z.* The marker colors indicate the ΔCCS values. P: phosphopeptides, uP: unphosphorylated peptides.

For further investigation, we focused on peptides in the extended subpopulation of triply charged species. The extended forms were visually classified based on their distributional shapes (Figure S1). In addition, to assess the phosphorylation position dependence of CCS changes, only Class I phosphopeptides^42^ (phosphosite localization probability > 0.75) were considered, leaving 277 pairs of phosphopeptides and unphosphorylated peptides. These pairs were used to examine the relationship between ΔCCS and the relative position of the phosphorylated amino acid residue within the peptide. It appeared that phosphorylation in the terminal region of the peptide is more likely to cause conformational compression, although the tendency is not large (Figure 6). Although there is insufficient molecular structure information to interpret the CCS data properly, the observations might be explained in terms of intramolecular interactions involving the phosphate groups. Phosphorylated residues are thought to be involved in multiple intramolecular interactions of peptide ions, and previous work^30,43^ on several doubly charged phosphopeptide ions has suggested that intramolecular salt bridges or ionic hydrogen bonds are formed between phosphate and basic groups in specific sequences.

**Figure 6.**
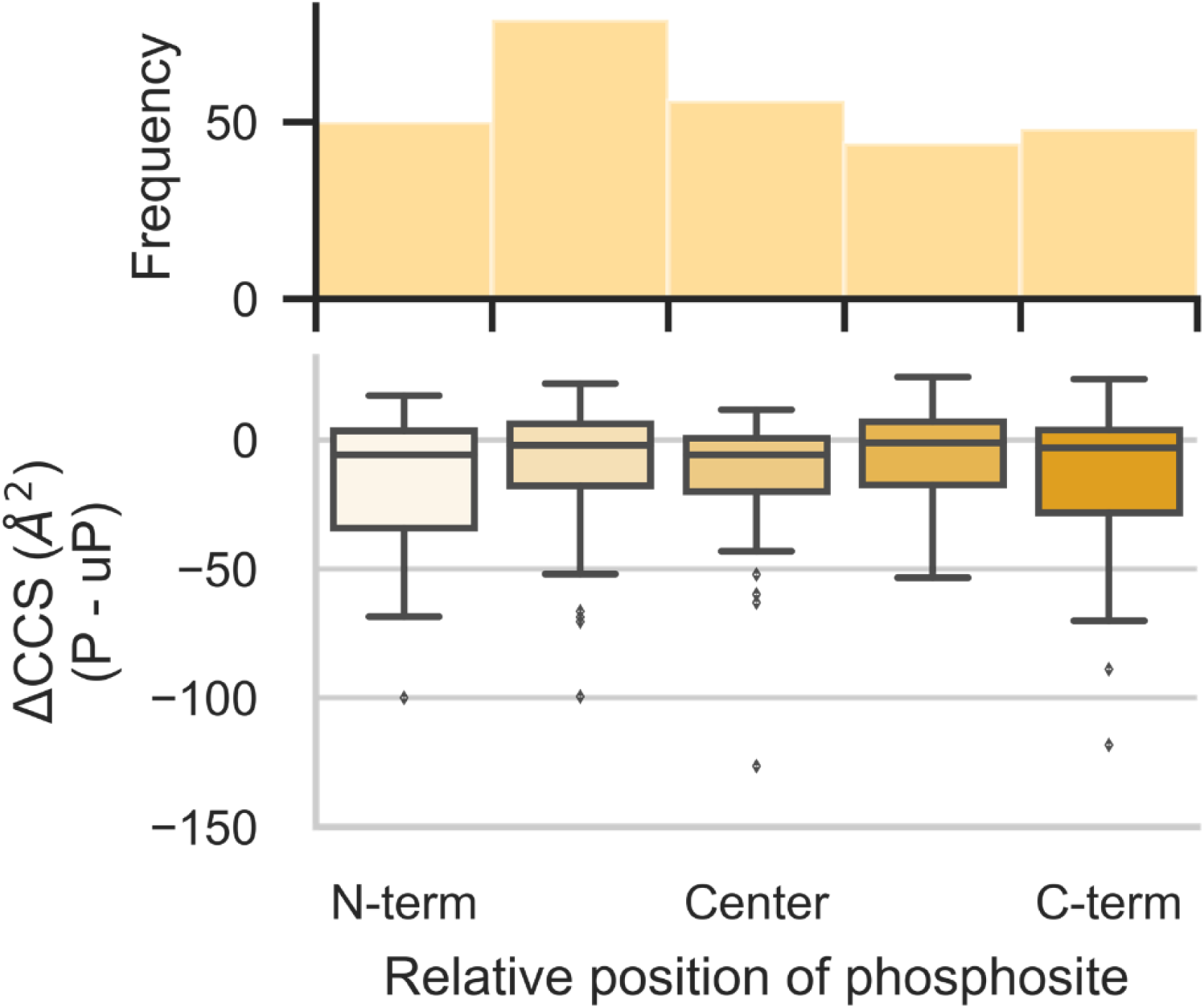
Relationship between phosphosite position in peptide and ΔCCS. 277 peptide pairs were categorized according to their relative phosphosite position. Box plots show the upper quartile, median, and lower quartile for the ΔCCS values for each subset. Outliers were identified using box-plot statistics (threshold: 1.5 x the interquartile range (IQR)).

It is possible that the phosphate group and its distal basic group may interact to compress the peptide structure, as shown in previous studies^30,43^. Another possibility is that the phosphate group may interact with its proximal basic group to disrupt the balance of positive charges in the peptide sequence, which would lead to changes in the peptide structure^38,44,45^. In other words, terminal phosphorylation may alter the charge balance within the peptide, leading to structural change.

To further evaluate these possibilities, we investigated the effects of basic amino acids on ΔCCS. In the dataset of 277 triply charged pairs, we found that the number of basic groups (Lys, His, Arg, and N term) of the peptide was weakly negatively correlated to ΔCCS (Figure 7A). This suggests that basic groups are involved in conformational compression upon phosphorylation. Considering that salt bridges or ionic hydrogen bonds require additional basic groups that do not contribute to the overall charge state of the peptide, peptides with more basic groups are more likely to have these intramolecular interactions. Therefore, we further focused on 127 of 277 triply charged peptide pairs with more basic groups than their charge state. These peptides would have basic groups that do not contribute to the total number of charges. Considering that the phosphate group in the terminal region of the peptide always has a proximal basic group derived from an N-terminal amine group or a C-terminal Lys or Arg, we examined the relationship between the ΔCCS and the distance between the phosphosite and its closest basic group, but no specific correlation was observed (Figure 7B). Next, we considered that the distance between other basic groups and the phosphorylation site may be important for the structural change of the peptide upon phosphorylation, and indeed, we found that the relative positional distance between the phosphosite and its second nearest basic group was weakly negatively correlated with ΔCCS (Figure 7C). These results can be explained by either of the above-mentioned mechanisms. Phosphorylation negates the positive charge of the closest basic group, and because the next closest basic group (or positive charge) is relatively far from the closest basic group, this disrupts the balance of charge in the peptide and causes a change in structure. Alternatively, an intramolecular interaction between the phospho site and the basic group distal to it may result in compression of the peptide ion structure. In any case, the results obtained in this study suggest that the interaction between phosphate and basic groups contributes significantly to the structural compression of phosphopeptides.

**Figure 7.**
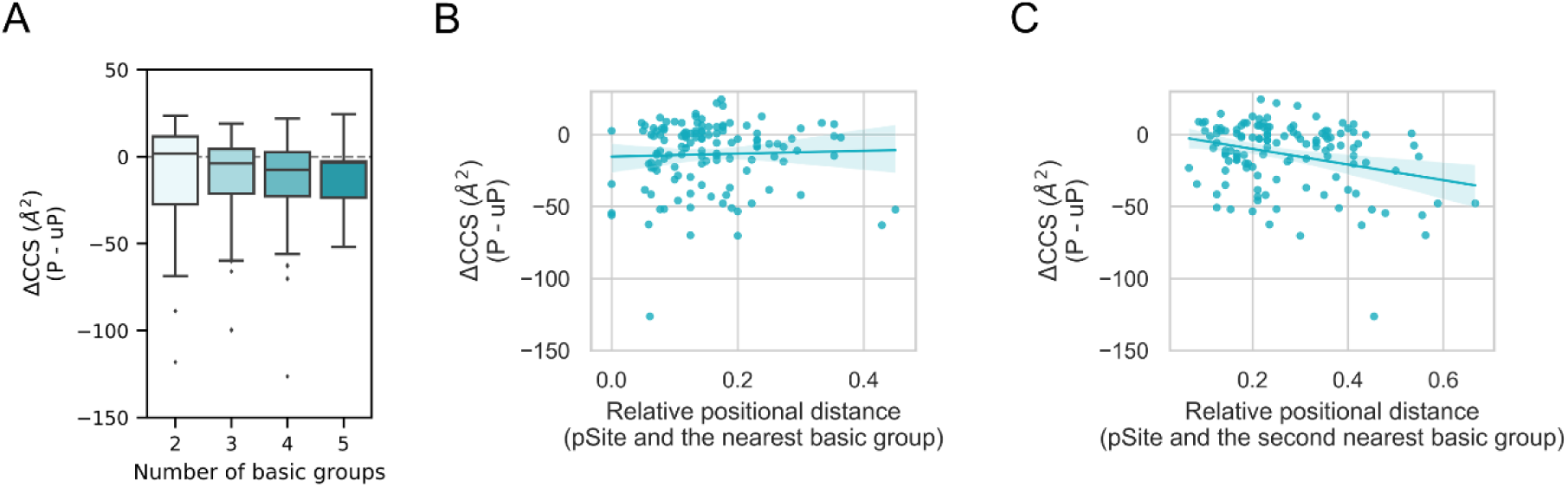
Relationship between basic groups in a peptide and ΔCCS. (A) 277 peptide pairs were categorized according to their number of basic groups. Box plots show the upper quartile, median, and lower quartile for the ΔCCS values in each subset. Outliers were identified using box-plot statistics (threshold: 1.5 x the interquartile range (IQR)). Dashed lines represent the ΔCCS value of zero. P: phosphopeptides, uP: unphosphorylated peptides. (B, C) ΔCCS values of 127 triply charged peptide pairs are plotted as a function of the relative positional distance between their phosphosite (pSite) and its nearest basic group. The relative positional distance was calculated by dividing the positional distance between the phosphosites and the basic groups by the length of the peptide sequence. The line represents a linear regression, and the shading indicates the 95% confidence interval.

## Conclusion

In this study, the CCS values of phosphopeptides and the corresponding unmodified counterparts were systematically profiled using TIMS. In agreement with previous reports, phosphorylation mainly results in compaction of peptide conformations. We also found that the CCS values of triply and quadruply charged ions are more strongly affected by phosphorylation, which could be explained at least in part by conformational changes of these larger peptides. The most prominent CCS changes were seen for triply charged peptides with extended structures and a larger number of basic groups, presumably reflecting intramolecular interactions between the phosphate group and the basic groups. We believe these findings will improve the CCS prediction of phosphopeptides.

## Acknowledgements

We would like to thank Ryo Kajita (Bruker Japan K.K.) for technical support and members of the Department of Molecular & Cellular BioAnalysis for fruitful discussions. This work was supported by JST Strategic Basic Research Program CREST (18070870), AMED Advanced Research and Development Programs for Medical Innovation CREST (18068699) and JSPS Grants-in-Aid for Scientific Research 17H03605.

## References

[1] Hunter T. Signaling—2000 and Beyond. Cell. 2000, 100, 113–127.

[2] Pawson T, Scott JD. Signaling through scaffold, anchoring, and adaptor proteins. Science. 1997, 278, 2075–2080.

[3] Mann M, Jensen ON. Proteomic analysis of post-translational modifications. Nat Biotechnol. 2003, 21, 255–261.

[4] Ishihama Y. Analytical Platforms for Mass Spectrometry-Based Proteomics. CHROMATOGRAPHY. 2019, 40, 89–97.

[5] Pasa-Tolić L, Masselon C, Barry RC, Shen Y, Smith RD. Proteomic analyses using an accurate mass and time tag strategy. Biotechniques. 2004, 37, 621-624, 626-633, 636 passim.

[6] Krokhin OV, Ying S, Cortens JP, et al. Use of peptide retention time prediction for protein identification by off-line reversed-phase HPLC-MALDI MS/MS. Anal Chem. 2006, 78, 6265–6269.

[7] Spicer V, Yamchuk A, Cortens J, et al. Sequence-specific retention calculator. A family of peptide retention time prediction algorithms in reversed-phase HPLC: applicability to various chromatographic conditions and columns. Anal Chem. 2007, 79, 8762–8768.

[8] Ma C, Ren Y, Yang J, Ren Z, Yang H, Liu S. Improved Peptide Retention Time Prediction in Liquid Chromatography through Deep Learning. Anal Chem. 2018, 90, 10881–10888.

[9] Moruz L, Tomazela D, Käll L. Training, selection, and robust calibration of retention time models for targeted proteomics. J Proteome Res. 2010, 9, 5209–5216.

[10] Gessulat S, Schmidt T, Zolg DP, et al. Prosit: proteome-wide prediction of peptide tandem mass spectra by deep learning. Nat Methods. 2019, 16, 509–518.

[11] Takahashi C, Yazaki T, Sugiyama N, Ishihama Y. Selected Reaction Monitoring of Kinase Activity-Targeted Phosphopeptides. CHROMATOGRAPHY. 2019, 40, 39–47.

[12] Schmidlin T, Debets DO, van Gelder CAGH, et al. High-Throughput Assessment of Kinome-wide Activation States. Cell Syst. 2019, 9, 366-374.e5.

[13] Ludwig C, Gillet L, Rosenberger G, Amon S, Collins BC, Aebersold R. Data-independent acquisition-based SWATH-MS for quantitative proteomics: a tutorial. Mol Syst Biol. 2018, 14, e8126.

[14] Kim J, Petritis K, Shen Y, Camp DG 2nd, Moore RJ, Smith RD. Phosphopeptide elution times in reversed-phase liquid chromatography. J Chromatogr A. 2007, 1172, 9–18.

[15] Perlova TY, Goloborodko AA, Margolin Y, et al. Retention time prediction using the model of liquid chromatography of biomacromolecules at critical conditions in LC-MS phosphopeptide analysis. Proteomics. 2010, 10, 3458–3468.

[16] Marx H, Lemeer S, Schliep JE, et al. A large synthetic peptide and phosphopeptide reference library for mass spectrometry–based proteomics. Nat Biotechnol. 2013, 31, 557–564.

[17] Ogata K, Krokhin OV, Ishihama Y. Retention Order Reversal of Phosphorylated and Unphosphorylated Peptides in Reversed-Phase LC/MS. Analytical Sciences. 2018, 34, 1037–1041.

[18] Meier F, Brunner A-D, Koch S, et al. Online Parallel Accumulation-Serial Fragmentation (PASEF) with a Novel Trapped Ion Mobility Mass Spectrometer. Mol Cell Proteomics. 2018, 17, 2534–2545.

[19] Myung S, Lee YJ, Moon MH, et al. Development of High-Sensitivity Ion Trap Ion Mobility Spectrometry Time-of-Flight Techniques: A High-Throughput Nano-LC-IMS-TOF Separation of Peptides Arising from a Drosophila Protein Extract. Anal Chem. 2003, 75, 5137–5145.

[20] Bonneil E, Pfammatter S, Thibault P. Enhancement of mass spectrometry performance for proteomic analyses using high-field asymmetric waveform ion mobility spectrometry (FAIMS). J Mass Spectrom. 2015, 50, 1181–1195.

[21] Bekker-Jensen DB, Martínez-Val A, Steigerwald S, et al. A Compact Quadrupole-Orbitrap Mass Spectrometer with FAIMS Interface Improves Proteome Coverage in Short LC Gradients. Mol Cell Proteomics. 2020, 19, 716–729.

[22] Ogata K, Ishihama Y. Extending the Separation Space with Trapped Ion Mobility Spectrometry Improves the Accuracy of Isobaric Tag-Based Quantitation in Proteomic LC/MS/MS. Anal Chem. 2020, 0c01695.

[23] Thalassinos K, Grabenauer M, Slade SE, Hilton GR, Bowers MT, Scrivens JH. Characterization of phosphorylated peptides using traveling wave-based and drift cell ion mobility mass spectrometry. Anal Chem. 2009, 81, 248–254.

[24] Glover MS, Dilger JM, Acton MD, Arnold RJ, Radivojac P, Clemmer DE. Examining the Influence of Phosphorylation on Peptide Ion Structure by Ion Mobility Spectrometry-Mass Spectrometry. J Am Soc Mass Spectrom. 2016, 27, 786–794.

[25] Ruotolo BT, Verbeck GF 4th, Thomson LM, Woods AS, Gillig KJ, Russell DH. Distinguishing between phosphorylated and nonphosphorylated peptides with ion mobility-mass spectrometry. J Proteome Res. 2002, 1, 303–306.

[26] Ruotolo BT, Gillig KJ, Woods AS, et al. Analysis of phosphorylated peptides by ion mobility-mass spectrometry. Anal Chem. 2004, 76, 6727–6733.

[27] Valentine SJ, Ewing MA, Dilger JM, et al. Using Ion Mobility Data to Improve Peptide Identification: Intrinsic Amino Acid Size Parameters. J Proteome Res. 2011, 10, 2318–2329.

[28] Valentine SJ, Counterman AE, Hoaglund-Hyzer CS, Clemmer DE. Intrinsic Amino Acid Size Parameters from a Series of 113 Lysine-Terminated Tryptic Digest Peptide Ions. The J Phys Chem B. 1999, 103, 1203–1207.

[29] Kaszycki JL, Shvartsburg AA. A Priori Intrinsic PTM Size Parameters for Predicting the Ion Mobilities of Modified Peptides. J Am Soc Mass Spectrom. 2017, 28, 294–302.

[30] Kim D, Pai P-J, Creese AJ, Jones AW, Russell DH, Cooper HJ. Probing the electron capture dissociation mass spectrometry of phosphopeptides with traveling wave ion mobility spectrometry and molecular dynamics simulations. J Am Soc Mass Spectrom. 2015, 26, 1004–1013.

[31] Masuda T, Tomita M, Ishihama Y. Phase transfer surfactant-aided trypsin digestion for membrane proteome analysis. J Proteome Res. 2008, 7, 731–740.

[32] Rappsilber J, Mann M, Ishihama Y. Protocol for micro-purification, enrichment, pre-fractionation and storage of peptides for proteomics using StageTips. Nat Protoc. 2007, 2, 1896–1906.

[33] Sugiyama N, Masuda T, Shinoda K, Nakamura A, Tomita M, Ishihama Y. Phosphopeptide Enrichment by Aliphatic Hydroxy Acid-modified Metal Oxide Chromatography for Nano-LC-MS/MS in Proteomics Applications. Mol Cell Proteomics. 2007, 6, 1103–1109.

[34] Prianichnikov N, Koch H, Koch S, et al. MaxQuant software for ion mobility enhanced shotgun proteomics. Mol Cell Proteomics. 2020, 001720.

[35] Cox J, Mann M. MaxQuant enables high peptide identification rates, individualized p.p.b.-range mass accuracies and proteome-wide protein quantification. Nat Biotechnol. 2008, 26, 1367–1372.

[36] Okuda S, Watanabe Y, Moriya Y, et al. jPOSTrepo: an international standard data repository for proteomes. Nucleic Acids Res. 2017, 45, D1107–D1111.

[37] Moriya Y, Kawano S, Okuda S, et al. The jPOST environment: an integrated proteomics data repository and database. Nucleic Acids Res. 2019, 47, D1218–D1224.

[38] Lietz CB, Yu Q, Li L. Large-scale collision cross-section profiling on a traveling wave ion mobility mass spectrometer. J Am Soc Mass Spectrom. 2014, 25, 2009–2019.

[39] Shah AR, Agarwal K, Baker ES, et al. Machine learning based prediction for peptide drift times in ion mobility spectrometry. Bioinformatics. 2010, 26, 1601–1607.

[40] Valentine SJ, Counterman AE, Clemmer DE. A database of 660 peptide ion cross sections: use of intrinsic size parameters for bona fide predictions of cross sections. J Am Soc Mass Spectrom. 1999, 10, 1188–1211.

[41] Taraszka JA, Counterman AE, Clemmer DE. Gas-phase separations of complex tryptic peptide mixtures. Fresenius J Anal Chem. 2001, 369, 234–245.

[42] Olsen JV, Blagoev B, Gnad F, et al. Global, in vivo, and site-specific phosphorylation dynamics in signaling networks. Cell. 2006, 127, 635–648.

[43] Simmonds AL, Lopez-Clavijo AF, Winn PJ, Russell DH, Styles IB, Cooper HJ. Structural Analysis of 14-3-3-ζ-Derived Phosphopeptides Using Electron Capture Dissociation Mass Spectrometry, Traveling Wave Ion Mobility Spectrometry, and Molecular Modeling. J Phys Chem B. 2020, 124, 461–469.

[44] Hudgins RR, Ratner MA, Jarrold MF. Design of Helices That Are Stable in Vacuo. J Am Chem Soc. 1998, 120, 12974–12975.

[45] Xiao C, Pérez LM, Russell DH. Effects of charge states, charge sites and side chain interactions on conformational preferences of a series of model peptide ions. Analyst. 2015, 140, 6933–6944.

