## Supplementary material for "Effect of Phosphorylation on the Collision Cross Sections of Peptide Ions in Ion Mobility Spectrometry": Figure S1

**Table of Contents**

**Figure S1. Visual classification of extended and compact forms of triply charged unphosphopeptides.**

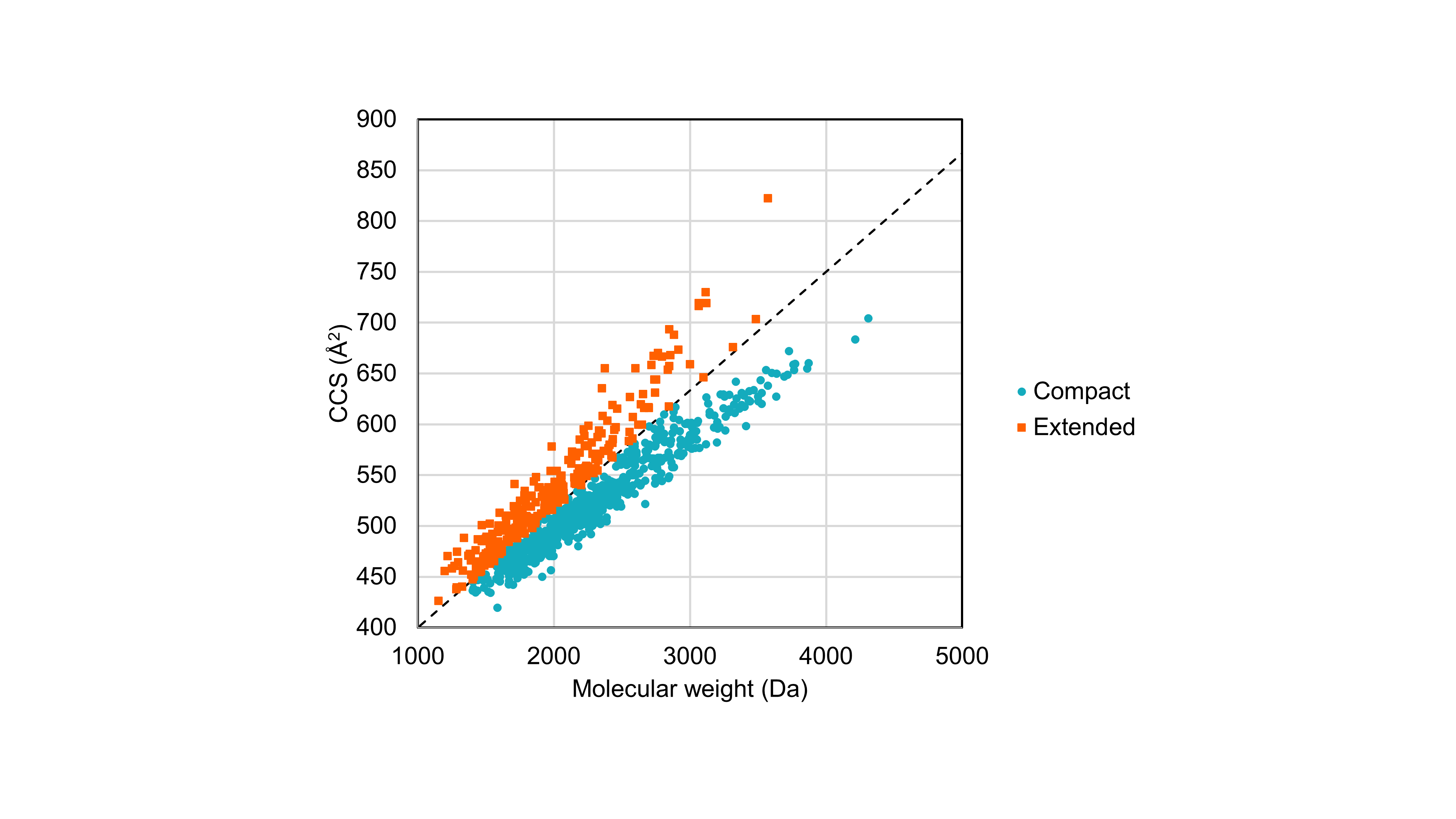

**Figure S1.**

Visual classification of extended and compact forms of triply charged unphosphopeptides. The dashed line represents the classification boundary.
